# Sequential functional differentiation of luminal epithelial cells regulated by maternal and embryonic signaling in the mouse endometrium

**DOI:** 10.1101/2022.04.30.490140

**Authors:** Hai-Quan Wang, Dong Li, Jingyu Liu, Yue Jiang, Ji-Dong Zhou, Zhi-Long Wang, Xin-Yi Tang, Yang Zhang, Xin Zhen, Zhi-Wen Cao, Xiao-Qiang Sheng, Chao-Fan Yang, Qiu-ling Yue, Li-jun Ding, Ya-li Hu, Zhi-Bin Hu, Chao-Jun Li, Gui-Jun Yan, Hai-Xiang Sun

**Affiliations:** Center for Reproductive Medicine and Obstetrics &Gynecology, Nanjing Drum Tower Hospital, The Affiliated Hospital of Nanjing University Medical School, Nanjing 210008, China; State Key Laboratory of Reproductive Medicine and China International Joint Research Center on Environment and Human Health, Center for Global Health, School of Public Health, Nanjing Medical University, Nanjing 211166, China; State Key Laboratory of Pharmaceutical Biotechnology, School of Life Sciences, Nanjing University, Nanjing 210023, China; Center for Molecular Reproductive Medicine, Nanjing University, Nanjing 210008, China

**Keywords:** luminal epithelial cells, functional differentiation, maternal-fetal crosstalk, scRNA-seq, thin endometrium

## Abstract

Embryo implantation requires temporospatial maternal-embryo dialog to regulate the interactions between an activated blastocyst and a receptive endometrium; however, the details of the related cell-specific coordination is largely unknown because of the cellular complexity and dynamic developmental processes of both the embryo and endometrium during peri-implantation. By comparing single-cell RNA (scRNA) data obtained from entire uteri of pregnant and pseudo-pregnant mice with embryos that were in the oviduct (dpc2.5 days post-coitum [dpc]) or had entered the uterus (after 3.5 dpc), we found a maternal estrogen and progesterone signaling-dependent functional differentiation process in which progesterone induced estrogen-responsive luminal epithelial cells to differentiate into two new types of epithelial cells: adhesion epithelial cells (AECs) and supporting epithelial cells (SECs) on 2.5 dpc. In addition to maternal signaling data, the uterine scRNA and corresponding embryo bulk sequencing data were obtained, and analyses revealed that embryonic Pdgfa and Efna3/4 signaling activated AECs and SECs, enhancing the attachment of embryos to the endometrium after implantation. Specifically, embryonic signaling induced additional transformation of AECs by preventing adjacent AECs, but not AECs away from the embryo in the implantation chamber, from undergoing apoptosis. Furthermore, we demonstrated that epithelial cell differentiation and related regulatory signaling were largely conserved in humans and mice. In the human endometrium, developmental defects of SOX9-positive epithelial cells similar to luminal epithelial cells were related to thin endometrium. Our work provides comprehensive and systematic information on endometrial luminal epithelial cell development directed by maternal and embryonic signaling, which is important for endometrial receptivity and embryo implantation.

## Introduction

Successful embryo implantation requires precise interactions between an activated blastocyst and receptive endometrium, and these interactions are profoundly influenced by a variety of temporal and spatial factors, especially crosstalk between embryogenic and maternal signaling components. Orchestrated by embryonic and maternal signaling, the endometrium becomes a suitable microenvironment for embryo implantation, which is known as “receptivity”, through a series of sequential cellular and molecular events^[1, 2]^. Thus, interactions between signaling molecules are critical for receptivity establishment, comprising the major limiting steps in mammalian reproduction. Indeed, abnormal alteration of signaling pathways may lead to pathological conditions and a low embryo implantation rate in humans^[3, 4]^.

During the establishment of receptivity, the morphology and cell function of the endometrium are altered to favor blastocyst implantation^[5]^, which is thought to be regulated by maternal ovarian steroid hormones. Sequential stimulation of estrogen and progesterone initiates changes in uterine structure and function that allow a blastocyst to attach and initiate the implantation process. In humans, estrogen induces the proliferation of epithelial, stromal, and vascular endothelial cells^[6]^. Then, progesterone secreted from the corpus luteum after ovulation promotes the endometrial transition from a proliferative to the secretory state, triggering the first step in establishing receptivity^[7]^.

At the same time, embryo-derived signals also affect embryo implantation. After entering the uterine cavity, the embryo secretes various signaling molecules, including growth factors, cytokines, and metabolites, before making physical contact with the luminal epithelium, which causes dramatic changes at the molecular and cellular levels^[8, 9]^. hCG is one of the earliest signaling molecules secreted by an embryo, and hCG is targeted to endometrial stromal cells to increase the expression of LIF and VEGF and further establishes and maintains pregnancy^[10]^. HB-EGF expression has been shown to be upregulated during blastocyst activation and induces the expression of Hb-egf in uterine cells surrounding blastocysts and increases vascular permeability^[11, 12]^. Blastocysts rapidly release unidentified cell-permeable lipids that activate uterine FAAH lipids and prevent abnormal endocannabinoid signaling, which would adversely affect preimplantation development and blastocyst implantation^[13]^. Embryo-derived signals can modulate the immune microenvironment of the maternal-fetal interface; for example, chemokines such as CXCL12 can recruit natural killer cells into the decidua^[14]^. Although we have identified some of the changes in the activities of endometrial cells during peri-implantation, how temporal and cell-specific dynamic alterations are sequentially regulated by maternal and embryogenic signaling is largely unknown because of the cellular complexity and dynamic changes in both the embryo and endometrium.

To explore the coordinated regulation of maternal and embryogenic signaling and their effects on the dynamic changes of endometrial cells during peri-implantation, we collected mouse uteri on 2.5, 3.5, 4.0 and 4.5 days post-coitum (dpc) and days pseudo-pregnant (dpp) for use in single-cell RNA sequencing (scRNA-seq) and embryos from the mouse uterus at 3.5 and 4.0 dpc for use in bulk RNA-seq. These samples were thus harvested at four time points from the time of endometrium preparation to embryo implantation. Since mouse embryos are still in the oviduct tube on 2.5 dpc, at this point the endometrium is regulated solely through maternal signaling. When embryos enter the uterus at 3.5 dpc or later, the endometrium receives signals from both the mother and embryo, which can be distinguished by comparing the pregnant mouse uterine data with the pseudo-pregnant uterine data.

Because of advances in scRNA-seq technology^[15]^, we identified the sequential differentiation of endometrial luminal epithelial cells associated with embryo implantation, which were controlled by maternal steroid hormone at 2.5 dpc. Through bioinformatics analyses, we identified cell communication between embryos and luminal epithelial cells from 3.5 to 4.5 dpc and identified embryo-derived peptide signaling that regulated blastocyst adhesion to luminal epithelial cells and promoted embryonic development.

## Materials and Methods

### Mice

All mouse experiments in this study were approved by the Institutional Animal Care and Use Committee of Nanjing Drum Tower Hospital (SYXK 2019-0059). Eight weeks old ICR female and male mice were purchased from Nanjing Medical University (Nanjing, China) and maintained in the Animal Laboratory Center of Nanjing Drum Tower Hospital (Nanjing, China) with a 12/12 h light/dark cycle (lights off at 1900 hours) with food and water available ad libitum.

### Uterine tissue collection

#### Normal pregnant mice

Eight weeks old ICR female mice were mated with fertile males. The morning that a vaginal plug was observed was termed day 0.5 of the pregnancy. The uterine tissues from the mice were collected on days 0.5, 1.5, 2.5, 3.5, 4.0 and 4.5 of pregnancy, divided into 0.5-1-cm sized samples, rapidly frozen in liquid nitrogen until they were stiff and preserved at -80°C.

#### Ovariectomy and hormone induction models

Eight weeks old ICR female mice were rested for 2 weeks after ovariectomy. Then, 50 µL of sesame oil containing 100 ng of E2 was injected subcutaneously for 3 consecutive days. The mice were allowed to rest for another 2 days, and then 50 µl of sesame oil containing 1 mg of P4 and 10 ng of E2 was injected subcutaneously for 2 consecutive days. The uterine tissues from the mice respectively at 2-weeks rest after ovariectomy, 6 h after receiving only the E2 injection, 2 days after receiving only the E2 injection, and 6 h after receiving the P4+E2 injection were collected and prepared into 0.5-1-cm pieces, rapidly frozen in liquid nitrogen until they were stiff and preserved at -80°C.

### Bulk RNA-seq library preparation and data analysis

Total RNA was isolated and used for RNA-seq analyses, and a cDNA library was constructed by Beijing Genomics Institute using an Illumina HiSeq X platform (Shenzhen, China). High-quality reads were aligned to the mouse reference genome (mm10) using HISAT2. The expression levels of each gene were calculated using FeatureCounts software. A differentially expressed gene (DEG) analysis was performed using the limma package, and all graphics were drawn with the ggplot package in R software.

### 10X Genomics library preparation, sequencing, and acquisition of an expression matrix

An scRNA-seq library was constructed following the 10X Genomics scRNA-seq protocol. Briefly, suspensions containing ∼10K cells were diluted following the instrument manufacturer’s recommendations and mixed with buffer before being loaded into a 10× Chromium Controller using Chromium Single Cell 3’ v3 reagents. Each sequencing library was prepared following the manufacturer’s instructions. The resulting libraries were then sequenced on an Illumina NovaSeq 6000 (Illumina, San Diego).

Raw sequencing data were demultiplexed using the mkfastq application (Cell Ranger v1.2.1). Three types of fastq files were generated: I1 contained an 8-bp sample index; R1 contained a 26-bp (10-bp cell [BC] + 16-bp unique molecular identifier [UMI]) index; and R2 contained a 100-bp cDNA sequence. The Fastq files were then run with the cellranger count application (Cell Ranger v1.2.1) using default settings to perform alignment (using STAR v2.5.4a), filtering and cell barcode and UMI counting. The UMI count tables for each cell barcode were used for further analysis.

### Data processing, batch effect correction and cell identification

We used the Seurat package^[16]^ to perform further analyses. Specifically, the UMI-based count matrix was first read into R using the Read10X function, and cells in which the number of genes was fewer than 500 or in which the mitochondrial gene ratio was more than 15% were considered low-quality cells and were removed. Finally, the remaining cells were used for subsequent analyses.

The NormalizeData function with default parameters was used to normalize the data. Then, the mean-variable of each gene was calculated using the vst algorithm in the FindVariableFeatures function, and 2,000 genes that induced the greatest differences among cells, that is, highly variable genes (HVGs), were selected. The ScaleData function was used to standardize the data for subsequent dimension reduction analysis.

We used the selected HVGs in a principal component analysis (PCA), then selected the top 10 principal components for batch correction with the harmony algorithm^[17]^ and used uniform manifold approximation and projection (UMAP) for further dimension reduction. For the identification of all cells, we first applied the FindNeighbor function to build a network and then used the FindCluster function to perform unsupervised cell clustering. The FindMarkers function was then used, and genes that me the log2 fold change (FC)>1 and p.adj<0.001 criteria were markers and were used to annotate cell populations.

### Pseudotime analysis

We used the Monocle2 package^[18]^ to construct a cell differentiation trajectory. Specifically, the UMI-based count matrix was read into R, and a CellDataSet was constructed (in Monocle). The HVGs selected by Seurat were used as the genes to calculate pseudotime, and the DDRTree algorithm in the reduceDimension function was used to construct a cell differentiation trajectory.

### Transcription factor-related gene regulation subnetwork analysis

Regulatory network and regulon activity analyses were performed with a mouse (mm10) dataset using the SCENIC package^[19]^. The UMI-based count matrix was used as input to identify co-expression modules through the GRNBoost2 algorithm. Then, regulons were derived by identifying the direct-binding transcription factor (TF) targets while pruning other regulons based on motif enrichment around transcription start sites (TSSs) in cisTarget databases. Using AUCell, the regulon activity score was determined as the area under the recovery curve (AUC), and the regulons status as active/inactive was identified based on the predicted AUC default threshold for each regulon in all cells. Subsequently, using a binarize function, we obtained the binary regulon activity score, which was converted into an ‘‘ON/OFF’’ (i.e., 1/0) labels. Furthermore, we removed regulons that were not active in at least 1% of the cells. We used a cutoff of 1% because it closely approximates the smallest cluster of the cells in our analysis, allowing the identification of regulons that were active in rare cell groups.

### Cell–cell communication analysis

To identify and visualize cell–cell interactions, we employed an R package named CellChat^[20]^. In brief, we followed the official workflow, loaded the normalized counts into CellChat and applied the standard preprocessing steps, including those using the functions identifyOverExpressedGenes, identifyOverExpressedInteractions, and projectData with a standard parameter set. A total of 2,021 precompiled mouse ligand– receptor interactions were selectively used as a priori network information. We then calculated the potential ligand–receptor interactions between embryo and epithelial cells based on the functions computeCommunProb, computeCommunProbPathway, and aggregateNet using standard parameters.

### Gene enrichment analysis

We performed Gene Ontology (GO) term and Kyoto Encyclopedia of Genes and Genomes (KEGG) enrichment analyses using enrichGO and enrichKEGG in the ClusterProfiler package^[21]^. DEG lists were used as input, and the appropriate organism of interest was used as the background. Terms with corrected P values < 0.05 were significant.

We performed a gene set enrichment analysis (GSEA) using gseGO and gseKEGG in the ClusterProfiler package. In summary, we sorted the log2FC values of the two groups and regarded them as the input for the gseGO/gseKEGG analyses. Terms with a P value < 0.05 were significant. Finally, we performed a gene set variation analysis (GSVA) using the gsva function in the GSVA package^[22]^.

### RT–qPCR verification

Total RNA was extracted from cells after treatment with TRIzol (Ambion/Thermo Fisher Scientific) according to the manufacturer’s instructions. The concentration and purity of all RNA was tested after extraction, and the A260/A280 value was above 2.0. Reverse transcription was performed to generate cDNA using 5X All-In-One RT MasterMix (with AccuRT Ge5X All-In-One RT MasterMix and AccuRT Genomic DNA Removal Kit) (Applied Biological Materials) for RT–qPCR, which was performed according to the instructions provided with the ChamQ SYBR RT–qPCR Master Mix (without ROX) (Vazyme) and using the fluorescence reagent SYBR and a qTOWER^3^G touch instrument (Analytik Jena). Data were analysed using the 2^−ΔΔCt^ relative quantitative method in Microsoft Excel software. The primers are: 5’- GTGAAGTGTATCGGGGAGCC- 3’ and 5’- TCCCGTTCCTTCAGAAAGGC -3’ for Epha1; 5’- ATGGCCGTTCTTAGTTGGTG- 3’ and 5’- CGGACATCTAAGGGCATCAC -3’ for 18S rRNA; 18S rRNA served as an internal control.

### In situ hybridization

In brief, frozen sections (10 μm) of mouse uterine tissues were fixed in 4% paraformaldehyde (PFA) in phosphate-buffered saline (PBS) solution at room temperature for 20 min, treated with gradient solutions of 0.25% and 0.5% acetic anhydride for 5 min each, and a preheated denatured digoxin (DIG) probe was added, and then, the samples were covered with silanized glass and incubated overnight at 60°C. The sections were washed twice with hybrid wash solution at 60°C for 30 min each time and then cooled naturally to room temperature. The sections were washed successively with MABT solution at room temperature for 30 min and RNA solution at 37°C for 10 min. Then, 80 μl of 10 mg/ml RNase was added to the RNA solution, and the treatment was continued at 37°C for 30 min. Next, the sections were treated successively with fresh RNA solution at 37°C for 5 min and fresh MABT solution at room temperature for 5 min. After blocking, the sections were incubated overnight with anti-digoxin antibodies (1:200, Roche, Indianapolis, IN) bound to alkaline phosphatase at 4°C. After washing with MABT solution and NTM solution, the sections were treated with 0.5 mg/mL levamisole for 10 min. Using BCIP/NBT (Beyotime Biotechnology) as the substrate, the staining signal was visualized according to the manufacturer’s instructions. Subsequently, the sections were stained with 0.1% nuclear fast red solution (Phygene) and sealed.

### Immunohistochemistry staining

Tissue sections were immunostained overnight with primary antibodies against Tacstd2 (1:1000, Abcam) at 4°C. On the following day, the sections were incubated with a goat anti-rabbit secondary antibody (a rabbit ABC detection kit, ZSBio, Beijing, China) at 37°C for 30 min. Next, the sections were stained with 3,3’-diaminobenzidine (DAB) and counterstained with hematoxylin. Control sections were prepared concurrently with the experimental sections and treated with nonspecific rabbit IgG. Unspecific staining was not detected in the controls.

### Mouse embryo transplantation assay

The mice on 3.5 dpc were sacrificed by neck dislocation, and blastocysts were removed and placed in M2 for future use. In addition, after anesthesia, 10 μl of a 100 μM PDGFR inhibitor, imatinib (Selleck), or 30 μl of a 20 μM Epha1 short interfering RNA (siRNA) mixture was injected into one uterine horn of the mice on 2.5 dpp and the same amount of control fluid was injected into the other, and then blastocysts were transplanted into each side of the uterus. The specific siRNA target sequences against Epha1 were GGATGCAAAGAGACCTTCA (#1), CACCACATCTACCGTGCAA (#2) and CCACATACATTCTCAGAGT (#3). The sutured mice were placed in a warm chamber and then sent to the breeding room after they were fully mobile. After 48 h, the mice were completely anesthetized and injected with Chicago blue dye in the tail vein. The number of blastocyst implantation sites in the mice was counted.

### Statistical analysis

The data are expressed as the means ± SEM. All experiments were performed at least three times. GraphPad Prism software (version 9.0) was used to perform statistical analyses. Statistical analyses were performed with one-way ANOVA for experiments involving more than two groups. P values less than 0.05 were statistically significant.

## Results

### Endometrial luminal epithelial cells show dramatic transcriptomic changes in the uterine receptivity state

Uterine receptivity needs to be established at the best time and in the correct place, which involves both functional and morphological alterations of a variety of cell types, including epithelial cells, stromal cells, endothelial cells, and immune cells, in the endometrium. To explore the dynamic changes in these cell types during embryo implantation in the uteri in the receptivity state, we compared the single-cell transcriptome profiles isolated from the whole uteri of female mice on 4.0 dpc when embryos were undergoing implantation and 4.0 dpp without embryo stimulation. In the uteri of the mice on 4.0 dpc, we identified 14 clusters (Sfig. 1A), which were assigned cell identities based on the expression of known markers (Sfig. 1B, 1C). These clusters were grouped into six main cellular categories: (1) epithelial cells (luminal and glandular); (2) stromal cells (mature stromal cells, Clec3b-positive stromal cells, and Mki67-positive proliferative stromal cells); (3) muscle cells; (4) endothelial cells; (5) mesothelial cells and (6) immune cells. When comparing the cell ratio between the uteri obtained on 4.0 dpc and 4.0 dpp, it showed that there were more proliferative stromal cells and fewer B cells in embryo implanted uteri (Sfig. 1D). In addition to the cell number comparison, we further explored cell activity changes by comparing the transcriptome in the uteri between 4.0 dpc and 4.0 dpp. Epithelial cells, including luminal and glandular epithelial cells, exhibited the most significant alteration in DEGs (Sfig. 1E). The GO analysis showed that these DEGs participated in processes including epithelial cell proliferation, cell-substrate adhesion, and various metabolic processes, e.g., oxidative phosphorylation, collagen metabolism and steroid metabolism (Sfig. 1F). These results suggested that the activity of the endometrial epithelium, especially the luminal epithelium that is in contact with the embryo first, displayed the most dynamic transcriptomic changes in the presence of embryos in the receptivity state; however, no notable change in the size of the epithelial cells population was found.

### A single-cell map of the whole mouse uterus during the peri-implantation period

Our hypothesis of endometrial alteration during the peri-implantation period suggests precise coordination through temporal and spatial crosstalk of embryogenic and maternal signaling molecules. Therefore, we isolated cells from one side of the uterus and/or embryos from another side of the uterus on 2.5, 3.5, 4.0 and 4.5 dpc to explore the effects of maternal and embryonic factors on the endometrial transcriptome at single-cell resolution. The effect of maternal and embryonic signaling was distinguished by comparing the uteri of pregnant mice with pseudo-pregnant mice. In addition, the bulk transcriptome data of embryos were used to analyze the embryonic signaling to endometrial cells at various time points (Fig. 1A).

**Figure 1.**
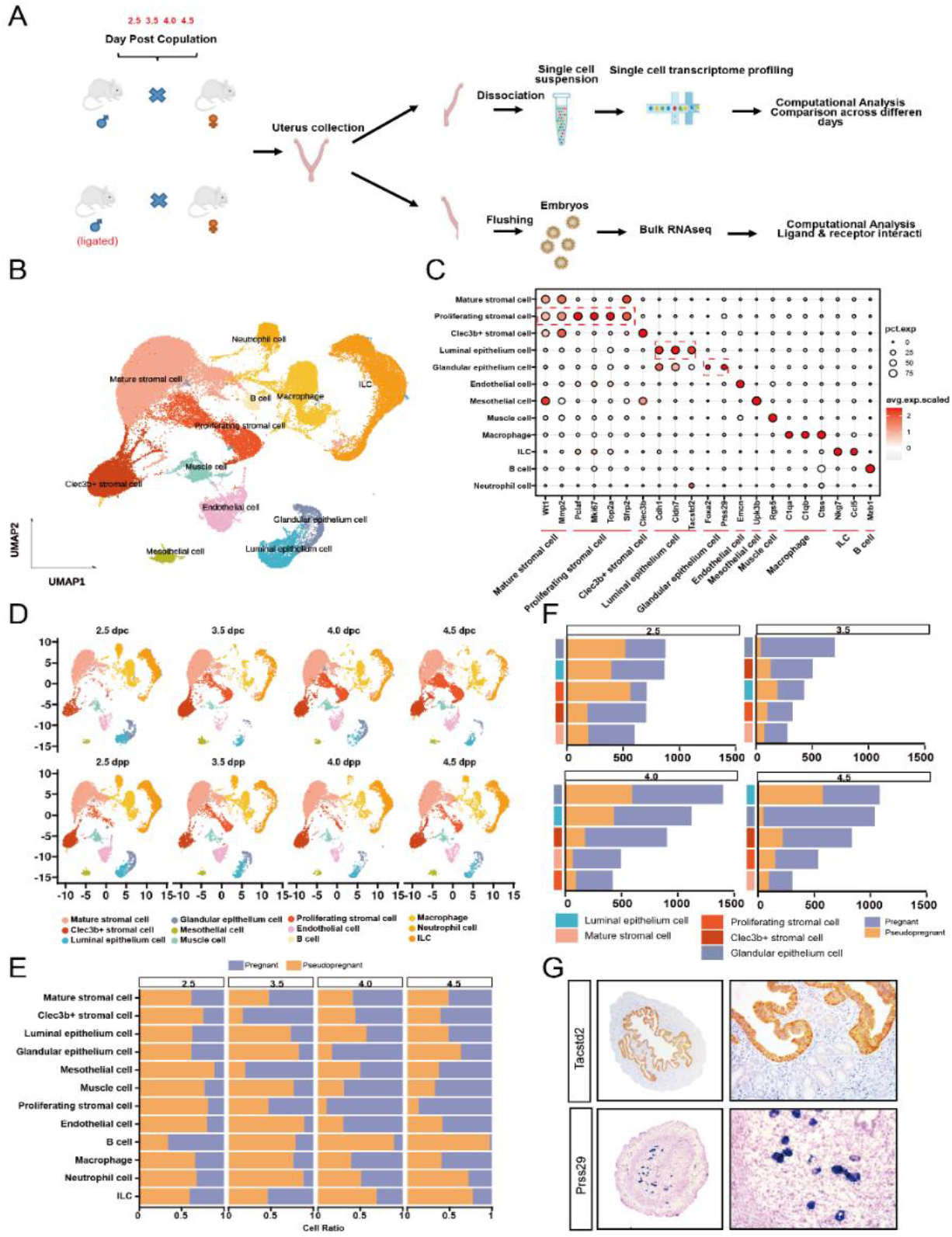
Single-cell profiling of the mouse uterus in pregnancy and pseudo-pregnancy. (A) Overview of the experimental design yielding scRNA-Seq and bulk RNA-Seq data from endometrium in pregnant or pseudo-pregnant mice. (B) UMAP projections of scRNA-seq data from endometrium in pregnant and pseudo-pregnant mice. (C) Dot plot showing log2-transformed expression of marker genes to identity cell types in endometrium. (D) UMAP of cells from endometrium of each mouse to check batch effect. (E) Bar plot showing relative proportion of cells in pregnant and pseudo-pregnant mice at each time point. (F) Bar plot showing the count of differential expressed genes in endometrial cell population between pregnant and pseudo-pregnant mice at each time point. (G) Immunohistochemistry of Tacstd2 and in situ hybridization of Prss29 expressing in uterus of mice.

For the cell lineage analysis of the uterus, 31647 cells passed standardized quality control (QC) filtering and were qualified for downstream analysis. We grouped the cells into 12 clusters based on the expression of known markers of different cell types (Fig. 1B, 1C). Batch effects were generally eliminated, as shown in Fig. 1D. We compared the cell ratio of different clusters in the pregnant and pseudo-pregnant mice at different time points and found significant differences in the cell ratio of multiple cell types (Fig. 1E). For example, the decrease in the ratio of B cells in pregnant uteri was dramatic from 2.5 to 4.5 dpc, in line with the establishment of immune tolerance as pregnancy progresses. In addition, the B cell ratio was continually increased in the pseudo- pregnant uteri. Interestingly, the ratio of proliferating stomal cells progressively increased in the pregnant uteri but decreased in the pseudo-pregnant uteri. Moreover, the ratio showed a different pattern, decreasing on 3.5 dpc when an embryo entered the uterus and decreasing after 4.0 dpc when an embryo begins to implant into the endometrium, which indicated that embryonic implantation regulated endometrial activity (Fig. 1E). This finding was confirmed by analyzing the DEG counts in the dominant cell types (luminal epithelial cells, mature stromal cells, proliferating stromal cells, Clec3b+ stromal cells and glandular epithelial cells) at different stages (Fig. 1F). The number of DEGs in all the dominant cell types was increased embryo attachment to the uterus (4.0 and 4.5 dpc), and the luminal and glandular epithelial cells exhibited the most significant changes. To distinguish luminal and glandular epithelial cells for further investigation, we identified Tacstd2 as a specific marker for luminal epithelial cells and Prss29 as a specific marker for glandular epithelial cells and verified their specificity through immunohistochemistry and in situ hybridization, respectively (Fig. 1G). Here, we describe a single-cell map of the whole mouse uterus during the peri-implantation period, revealing endometrial cell activity alterations corresponding to the embryo entering the uterus and embryo implantation into the endometrium. Among different uterine cell types, luminal epithelial cells first contact an embryo and play important roles in embryo implantation; therefore, we next focused on changes in the luminal epithelium during the peri-implantation period.

### Luminal epithelial cells functionally differentiate into adhesion epithelial cells and supporting epithelial cells in the peri-implantation period

To further explore the dynamic changes of luminal, we performed a pseudo-time analysis using the Monocle2 software package, and the trajectory suggested cell differentiation across the pseudo-time, from “Root” cells at pseudo-time (0) into two bifurcated subtypes, “branch1” and “branch2” (Fig. 2A, Sfig2A). The “Root” cells were mainly composed of cells on 2.5 dpc, with “branch1” cells emerging 2.5 dpc, and “branch2” appearing 3.5 dpc, suggesting that the differentiation of luminal epithelial cells is initiated on 2.5 dpc. The “Root” cells almost completely disappeared, having differentiated into the two branches on 3.5 dpc. A similar pseudo-time trajectory was observed in the pseudo-pregnant uteri (Sfig 2B). We calculated the Pearson correlation coefficients of the three subtypes of luminal epithelial cells in the pregnant and pseudo-pregnant uteri, and the results revealed significant similarity in transcriptome features (R^2^>0.95) (Fig. 2B), which suggested that luminal epithelial cell differentiation was not induced by embryo-derived signaling; in contrast, it was probably driven by maternal signaling.

**Figure 2.**
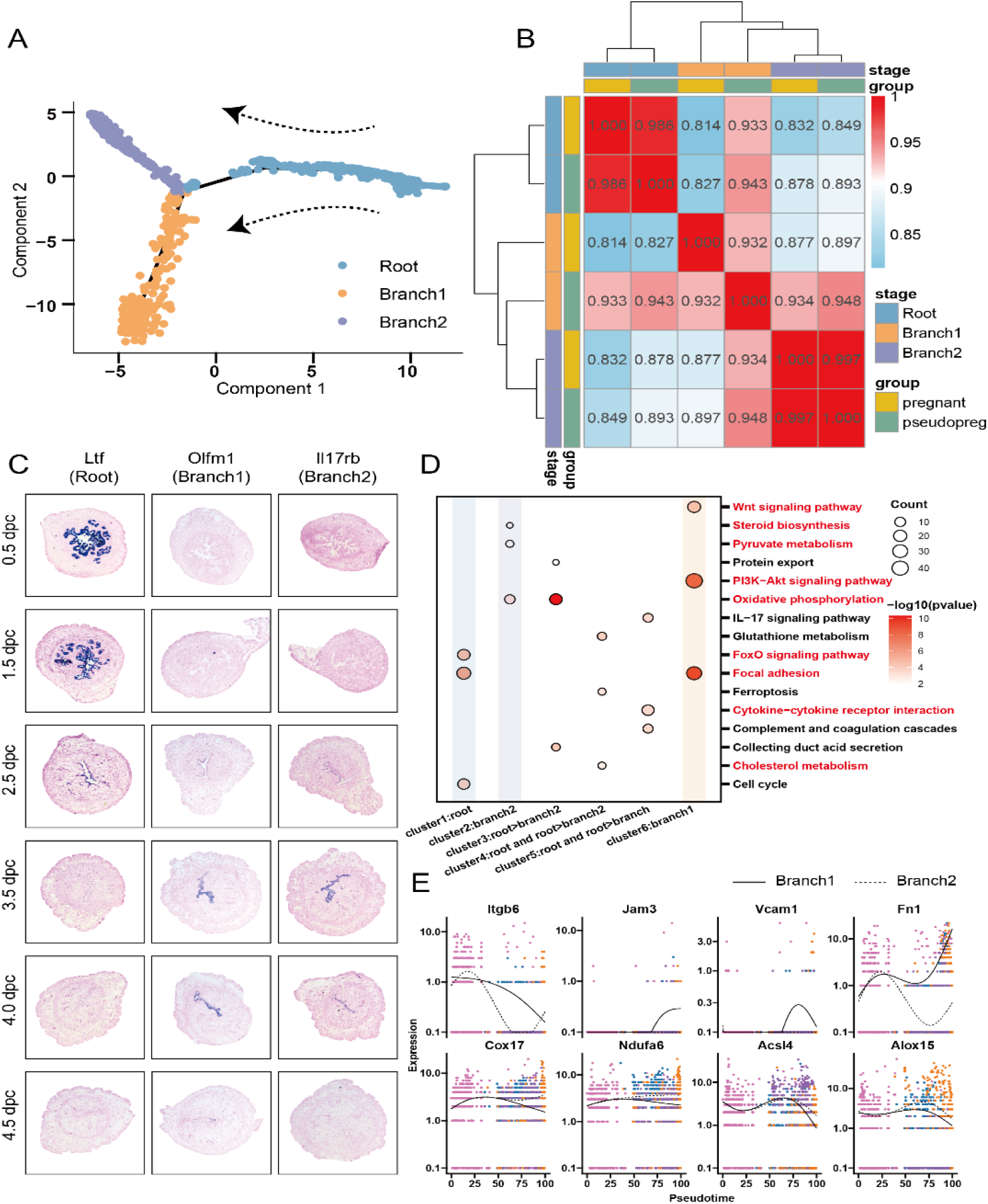
Functional differentiation of luminal “Root” epithelial cells during receptive establishment. (A) Developmental pseudo-time of endometrial luminal epithelial cells in pregnant mice. Arrows indicate the developmental order of these cells. (B) Heatmap showing pearson correlation coefficient of transcriptome of three types of endometrial luminal epithelial cells between pregnant and pseudo-pregnant mice. (C) In situ hybridization markers of luminal “Root” (Ltf), “Branch1” (Olfm1) and “Branch2” (Il17rb) in the uteri of mice on 0.5-4.5 dpc. (D) Dot plot showing KEGG enrichment results of clusters of genes that are differentially expressed across pseudo-time. Labels of hierarchical axis were according to cluster name from Sfig2D. Point size represented number of genes enriched in the pathways. The color bar represented -log_10_ -transformed P values of the pathways. (E) Expression levels (vertical axis) of key genes between Branch1 and Branch2 ordered in pseudo-time (cell colors were according to time points from Sfig2A).

The specific markers Ltf and Clu for “Root” cells, Atp6v0d2 and Olfm1 for “branch1” cells, and Il17rb and Nudt19 for “branch2” cells were identified to facilitate further experimental investigations (Sfig. 2C). In situ hybridization (ISH) verified that Ltf was specific to “Root” cells, while Olfm1 was specific to “branch1” cells, and Il17rb expression 3.5 and 4.0 dpc, but not 4.5 dpc, was specific to “branch2” cells (Fig. 2C). In addition, the ISH results confirmed that the “Root” cells began to differentiate on 2.5 dpc and were completely transformed by 3.5 dpc. “Branch1” cells appeared from 2.5 to 4.5 dpc, while “branch2” cells emerged starting 3.5 dpc, continually emerging until 4.5 dpc.

A heatmap analysis revealed the existence of three transient stages in association with “Root”, “branch1” and “branch2” cells, and these cells at all six stages exhibited significantly different transcriptomic features (Sfig. 2D). A KEGG analysis of the six gene sets showed that “branch1” cells highly expressed genes associated with “gap junctions”, “focal adhesions” and “extracellular matrix (ECM)-receptor interactions”, while “branch2” cells highly expressed genes related to “oxidative phosphorylation”, “pyruvate metabolism” and “thermogenesis”. “Root” cells expressed genes associated with the “cell cycle” (Fig. 2D), which was also proven by the GSVA scores obtained for all three types of epithelial cells (Sfig. 2E). A pseudo-time-related gene expression analysis of several key markers showed that “branch1” cells expressed higher levels of genes associated with intercellular adhesion (Itgb6, Jam3, and Vcam1) and ECM adhesion (Col1a2 and Fn1), while “branch2” cells expressed higher levels of genes related to metabolic activity (Cox17, Ndufa6, Atp5j, and Atp6v0e) and ferroptosis (Acs14 and Alox15) (Fig. 2E). Thus, we assumed that “branch1” cells probably directly adhere with embryos and thus were named adhesive epithelial cells (AECs). In contrast, “branch2” cells exhibited gene expression patterns indicative of metabolic support and therefore were named supporting epithelial cells (SECs). We confirmed their function in a subsequent experiment.

In conclusion, our results suggested a functional differentiation of luminal epithelial cell into AECs and SECs during receptivity establishment. Since differentiation was initiated on 2.5 dpc, when embryos are still in the oviduct, we anticipated that differentiation be induced by maternal not embryonic factors.

### Maternal estrogen and progestogen signaling is critical for the functional differentiation of luminal epithelial cells during receptivity establishment

To further elucidate the maternal factors that induce functional differentiation of the luminal epithelial cells, we constructed ovariectomized mouse models and mimicked the endocrinal states at the receptivity state by injecting estrogen and progestogen (Fig. 3A). We found that the appearance of “Root” luminal epithelial cells depended on stimulation by estrogen, which increased after coitus regardless of whether the mice were pregnant (Fig. 3B). The luminal “Root” epithelial cells were found up to 2 days after estrogen treatment was stopped, but the other groups of cells did not appear at this time (Fig. 3B). After injection of estrogen and progesterone (E+P stimulation at 19th days after ovariectomy), “root” luminal epithelial cells completely disappeared, and the other two groups of cells, the AECs and SECs, appeared on the luminal surface of the endometrium (Fig. 3B), which indicated that the functional differentiation of AECs and SECs was dependent on the stimulation of progesterone, which was increased during receptivity establishment. Thus, the sequential stimulation of the maternal factors estrogen and progestogen was critical for the functional differentiation of luminal epithelial cells during peri-implantation. Therefore, “Root” luminal epithelial cells were named estrogen-responsive epithelial cells.

**Figure 3.**
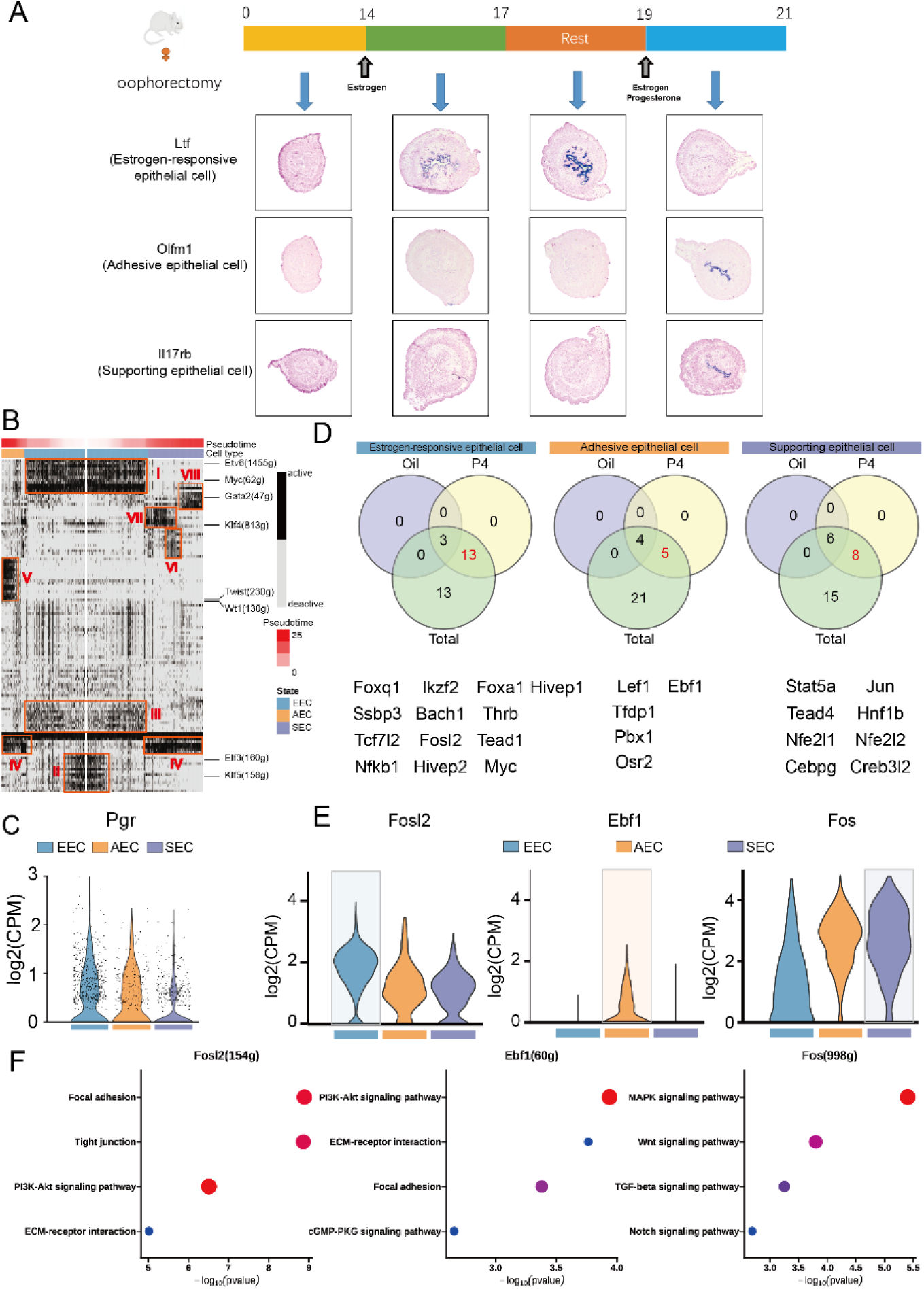
Functional differentiation of luminal “Root” epithelial cells was induced by maternal signals. (A) Model diagram of hormone model mice uteri samples collection and in situ hybridization markers of estrogen-responsive epithelial cells (Ltf), adhesive epithelial cells (Olfm1) and supporting epithelial cells (Il17rb) in the uteri of mice. (B) Heatmap showing activities of regulons in luminal epithelial cells according to SCENIC analysis. Black represented active status of regulons and grey represented deactive status of regulons. (C) Violin plot showing log_2_ -transformed expression of Pgr in three types of luminal epithelial cells. (D) Venn diagram showing relationship between TFs filtered from oil-treated and P4-treated mouse uterus and regulons derived from SCENIC analysis. Oil represented TFs filtered from oil-treated mouse uterus. P4 represented TFs filtered from progesterone-treated mouse uterus. Total represented regulons (TFs) derived from SCENIC analysis. (E) Violin plot showing log_2_ -transformed expression of key TFs in three types of luminal epithelial cells. (F) Dot plot showing KEGG enrichment results of target genes regulated by key TFs.

To determine the effect of progesterone on the functional differentiation of the luminal epithelial cells, we constructed TF subnetworks or ‘‘regulons’’ using SCENIC and three types of epithelial cells. On the basis of pseudo-time order, we found that sequential regulon activation in different epithelial cells followed distinct patterns (Fig. 3B, I-VIII). Notably, regulons such as Elf3 and Klf5 were activated in the epithelial cells and were subsequently deactivated during differentiation (Fig. 3B, I), while Etv6 and Myc continued to be activated until differentiation was completed (Fig. 3B, II). In AECs, regulons such as Twist1 and Wt1 were activated, and both are related to the epithelial-mesenchymal transition (EMT) (adhesion) (Fig. 3B, V). In SECs, Klf4 was activated first (Fig. 3B, VII), followed by Gata2 (Fig. 3B, VIII). Klf4 can increase the level of oxidative phosphorylation, indicating that Klf4 may be an important TF for increasing the SEC metabolism rate^[23]^. Furthermore, we found that all three types of epithelial cells expressed Pgr, a gene that encodes progesterone receptors (Fig. 3C). Performing a genome-wide PR interaction site analysis using chromatin immunoprecipitation and sequencing (ChIP-seq) data^[24]^ from the GSE34927 dataset based on mouse uteri treated with P4 and integrating SCENIC results, we found that 13 TFs in luminal “Root” epithelial cells, 5 TFs in AECs, and 8 TFs in SECs were bound by PGR (Fig. 3D). We then constructed a regulatory network with the TFs in all three epithelial cell types and found that the key TF in EECs was Fosl2 for luminal “Root” epithelial cells, was Ebf1 in AECs and was Fos in SECs, which might be important for the function of all three types of epithelial cells (Fig. 3E, Sfig. 3A). A KEGG enrichment analysis confirmed the functional relationships of the target genes of these TFs, which were aligned with AEC and SEC functions: the downstream genes of Fosl2 were involved in focal adhesion, tight junctions and ECM-receptor interactions; the target genes of Ebf1 were the PI3K-Akt signaling pathway, ECM-receptor interactions and focal adhesion; and Fos was related the MAPK, WNT and TGF-beta signaling pathways (Fig. 3F). These results suggested that progesterone might direct luminal “Root” epithelial cell differentiation by activating TF through the PGR in epithelial cells.

In summary, our data indicated that maternal estrogen was critical for the appearance and maintenance of luminal “Root” epithelial cells, while progestogen signaling was critical for the functional differentiation of luminal “Root” epithelial cells to AECs and SECs by activating the regulon through downstream TFs.

### Embryo-derived PDGF signaling regulates AEC activity and mediates embryo-endometrial interactions during implantation

We showed that luminal “Root” epithelial cells to AEC and SEC differentiation was stimulated by maternal progesterone approximately on 2.5-3.5 dpc when embryos began to enter the uterus. Since the progesterone surge was not related to pregnancy, luminal “Root” epithelial cell to AEC and SEC differentiation was also observed in pseudo-pregnant mice. However, when we compared the transcriptome profiles of AECs on 3.5, 4.0 and 4.5 dpc with those of their pseudo-pregnant counterparts, the number of DEGs was significantly enhanced on 4.0 dpc, when embryos began to implant in uteri, compared with the number counted 3.5 or 4.5 dpc (Fig. 4A). This finding indicated that embryo-derived signaling induced the transcriptome to that consistent with AECs, which might have been important to establish receptivity and ensure embryo implantation. Of course, the maternal factors, including progesterone and endometrial microenvironment signaling molecules, were still expressed during peri-implantation. To dissect the embryo-derived signaling from maternal factors, for all time points (embryonic days [E] 3.5, 4.0 and 4.5) of our bulk embryo sequence data and single-cell sequence datasets, we obtained ligand–receptor pairs in which ligands were from embryos and the receptors were from AECs, as calculated by CellChat. Then, receptors that were expressed in pseudo-pregnant mice and did not differ from those expressed in pregnant mice were removed. The retained the ligand–receptor pairs, named embryo-to-AEC LRs, were assumed to be secreted from the embryo and received by AECs. Using this research strategy, we identified 18 LR pathways (Fig. 4B), among which the WNT, EGF and SPP1 signaling pathways have been reported to mediate embryo-endometrium crosstalk, while the EPHB, PDGF and PROS signaling pathways had not been previously reported. To determine the influence of embryo-derived signaling on AECs, DEGs between pregnant and pseudo-pregnant mice identified at each time point were considered potential embryo ligand-related DEGs.

**Figure 4.**
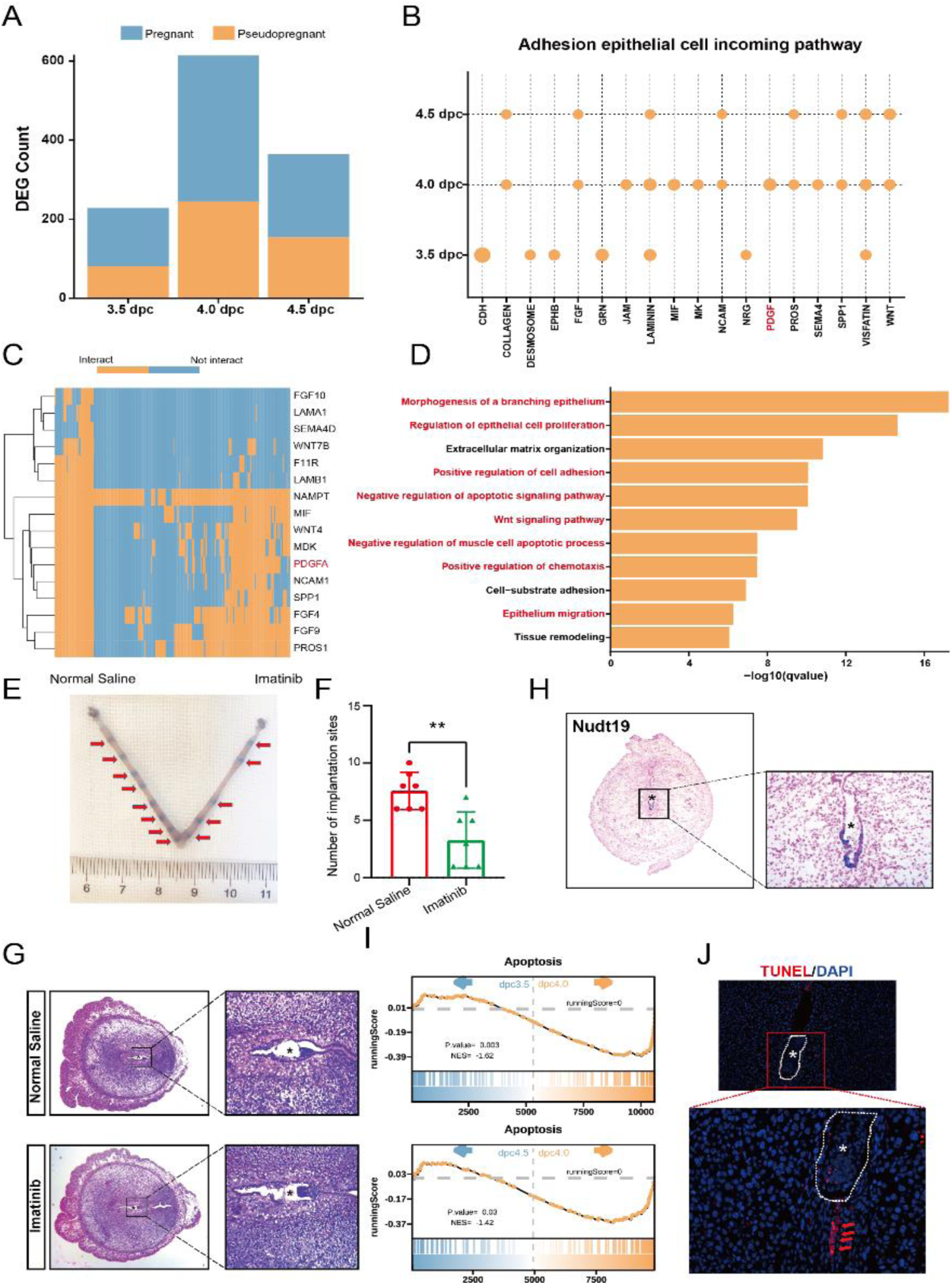
Embryo-derived signals regulated AEC activity and implantation. (A) Bar plot showing the number of DEGs in AECs between pregnant and pseudo-pregnant mouse at three time points respectively. (B) Dot plot showing which embryo-derived signals interacted with AEC at pathway levels. (C) Heatmap showing DEG of AEC between pregnant and pseudo-pregnant mouse uterus targeted by embryo-derived signals. “Interact” meant the signals could potentially regulated the target genes and “Not interact” meant the signals couldn’t potentially regulated the target genes. (D) Bar plot showing enriched GO terms of genes target by embryo-derived signals. (E) Representative uteri treated with normal saline (uterus on the left) or PDGF inhibitor imatinib (uterus on the right) on day 4.5 (n=6). (F) Number of embryo implantation sites in uteri treated with normal saline (uterus on the left) or PDGF inhibitor imatinib (uterus on the right) on day 4.5 (n=6). (G) Hematoxylin-eosin staining of mice uteri treated with normal saline (uterus on the left) or PDGF inhibitor imatinib (uterus on the right) on day 4.5. (H) In situ hybridization markers of adhesive epithelial cells (Nudt19) in the uteri of mice on 4.5 dpc. (I) GSEA plot of the KEGG apoptosis pathway for AECs between 3.5dpc and 4.0dpc (top) and between 4.5 dpc and 4.0dpc (bottom). (J) TUNEL staining of the uteri of mice on 5.5 dpc.

Then, we predicted the regulatory relationships between embryo-to-AEC LRs and embryo-related DEGs using the nichenetR package. We identified 16 ligands from 18 LR pairs that were also derived from embryos (Fig. 4C), including WNT7B and WNT4, as well as PDGFA, which had not been previously studied. Moreover, a GO enrichment analysis revealed that the embryo related DEGs were mainly involved in epithelial cell remodeling, including morphogenesis of branching epithelial cells and epithelial cell migration and adhesion, including ECM organization and the positive regulation of cell adhesion (Fig. 4D). Hence, we speculated that embryo-derived signaling regulates embryo-AEC adhesion. To verify our hypothesis, we injected the PDGFR antagonist Imatinib on 2.5 dpc and then transplanted embryos into mice. We found that the number of embryo implantation sites in the Imatinib-treated uteri was significantly reduced on 4.5 dpc, suggesting that the PDGF pathway plays a vital role in embryo implantation (Fig. 4E, 4F, Table 1). We also observed that the size of the implanted embryos was similar to that of the saline control group embryos (Fig. 4E). Hemoxylin and eosin (HE) staining of the implanted embryos indicated that embryonic development was not affected by PDGF treatment (Fig. 4G), although embryo implantation had been truly inhibited.

**Table 1.** Effects of PDGF signaling on embryo implantation.

| Variable | Groups |  | P value |
| --- | --- | --- | --- |
|  | Normal Saline | Imatinib |  |
| NO. of recipients | 6 | 6 | NS |
| No. of blastocysts transferred | 60 | 60 | NS |
| No. of implantation sites | 7.33±1.63 | 2.67±1.97 | 0.0022 |

Note: Values are presented as mean ± SD or number (percentage). NS = nonsignificant

After embryo attachment, the luminal epithelium invaginates to form an implantation chamber. Any abnormalities in the luminal epithelium during this process can result in compromised pregnancy outcomes. Herein, we showed that AECs were widely distributed in the endometrial lumen from 2.5 to 4.0 dpc and were retained until 4.5 dpc when AECs were located only around the implanted embryo in the implantation chamber (Fig. 2D). This finding indicated that AECs in other areas in the implantation chamber might have been eliminated after embryo implantation. The location of the AECs was confirmed by the expression of another AEC marker, Nudt19, which was found only near the implantation site (Fig. 4H). Based on a GSEA, we found that activation of the apoptosis pathway in the AECs was significantly upregulated on 4.0 dpc, when the embryo began to implant (Fig. 4I). In addition to embryo-derived signaling might preventing adjacent AECs from undergoing apoptosis, antiapoptotic processes, such as negative regulation of the apoptotic signaling pathway, responded to embryo-derived signaling in the AECs (Fig. 4D). We found many apoptotic signaling pathway components were expressed in the implantation chamber but not in the implantation site (Fig. 4J).

In conclusion, our results indicated that, although maternal signaling controls the functional differentiation of luminal “Root” epithelial cells into AECs, embryo-derived signaling affected AEC activity and fate during and after embryonic implantation in the endometrium. PDGFA, along with other embryo-derived factors, regulated AEC remodeling and adhesion to mediate the interaction of an embryo and the endometrial epithelium.

### Embryo-derived Efna/Epha signaling regulates SEC activities and affects both embryo implantation and development

Similar to that of the AECs, the transcriptome profile of the SECs from 3.5 to 4.5 dpc and that in of their pseudo-pregnant counterparts, indicated that the number of DEGs was significantly enhanced on both 4.0 and 4.5 dpc, which was after embryo implantation (Fig. 5A). Using the same strategy, we identified 9 LR pairs that were also ligands expressed by embryos and receptors expressed by SECs on 4.0 and 4.5 dpc (Fig. 5B), among which secreted WNT, EPHB, SPP1, VISFATIN, COLLAGEN and LAMNIN were directed to AECs, and BMP, FGF and EPHA were SEC-specific. Then, we identified 6 ligands produced by E4.0 embryos and 9 ligands produced by E4.5 embryos and predicted the interactions between ligands and embryo related DEGs in SECs using the nichenetR package. The ligands included Lamc2 and Lamb1, as well as Efna3 and Efna4, which had not been previously reported (Fig. 5C). Furthermore, a GO enrichment analysis revealed that these regulated genes were involved in protein phosphorylation, regulation of the apoptotic signaling pathways and positive regulation of catabolic processes on both 4.0 and 4.5 dpc (Fig. 5D). For verification, we knocked down Epha1 expression, which is the receptor for Efna3 and Efna4, in the SECs. Epha1 siRNA mixture was injected into one side of the uterus to knockdown its expression and control siRNA was injected on the other side on 2.5 dpc, and then both sides were implanted with the same number of embryos. We found that the number of embryos implantation sites in the Epha1 siRNA-treated uterus was significantly reduced on 4.5 dpc, suggesting that the Epha1 pathway plays a significant role in embryo implantation (Fig. 5E, 5F, Table 2). We also observed that the size of the implanted embryo was similar to that of the control group (Fig. 5E). HE examination of the implanted embryos indicated that embryonic development was not affected by Epha1 knockdown (Fig. 5G), although embryo implantation had been truly inhibited.

**Table 2.** Effects of Efna/Epha signaling on embryo implantation.

| Variable | Groups |  | P value |
| --- | --- | --- | --- |
|  | siNC | siEpha1 |  |
| NO. of recipients | 5 | 5 | NS |
| No. of blastocysts transferred | 49 | 49 | NS |
| No. of implantation sites | 6.50±1.64 | 1.60±1.86 | 0.0008 |

Note: Values are presented as mean ± SD or number (percentage). NS = nonsignificant

**Figure 5.**
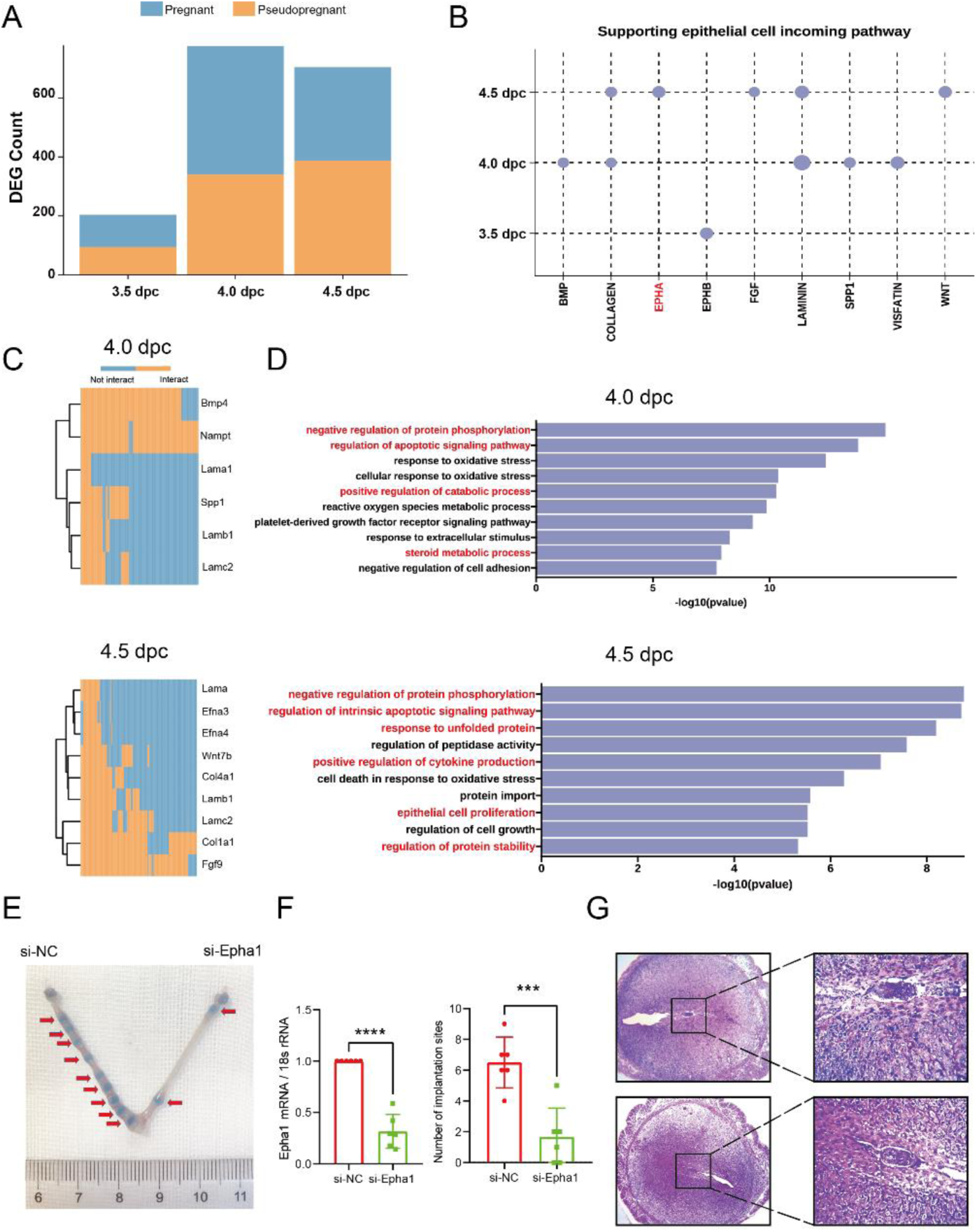
Embryo-derived signals regulated SEC activity and implantation. (A) Bar plot showing the number of DEGs in SEC between pregnant and pseudo-pregnant mouse at three time points respectively. (B) Dot plot showing which embryo-derived signals interacted with SEC at pathway levels. (C) Heatmap showing DEG of SEC between pregnant and pseudopregnant mouse uterus targeted by embryo-derived signals at 4.0 dpc and 4.5 dpc. “Interact” meant the signals could potentially regulated the target genes and “Not interact” meant the signals couldn’t potentially regulated the target genes. (D) Bar plot showing enriched GO terms of genes target by embryo-derived signals at 4.0 dpc and 4.5 dpc. (E) Representative uteri treated with NC siRNA (uterus on the left) or Epha1 siRNA mixture (uterus on the right) on day 4.5 (n=6). (F) mRNA expression levels of Epha1 in siNC and siEpha1 groups examined by RT-qPCR (left) (n=6) and number of embryo implantation sites in uteri treated with NC siRNA (uterus on the left) and Epha1 siRNA mixture (uterus on the right) on day 4.5 (right) (n=6). (G) Hematoxylin-eosin staining of mice uteri treated with NC siRNA (uterus on the left) or Epha1 siRNA mixture (uterus on the right) on day 4.5.

All these results indicated that embryo-derived signaling affected SEC activity, like its effect in AECs. Moreover, Epha1 regulated embryo implantation.

### Both luminal epithelial cell differentiation and directed signaling are conserved in humans and mice

Due to the menstrual cycle, human endometrial luminal epithelial cells are always in periodic proliferation or differentiation states. Four groups of epithelial cells are included in the human endometrium GSE111976 dataset: glandular epithelial cells, unciliated luminal epithelial cells, ciliated luminal epithelial cells, and Sox9-positive luminal epithelial cells ^[25]^. Using Monocle2, we constructed the trajectory of the differentiation of human luminal epithelial cells, and we found that SOX9-positive epithelial cells differentiated into two groups, unciliated luminal epithelial cells and ciliated luminal epithelial cells, from the proliferative stage through the secretory stage, similar to that in the mice in which luminal “root” epithelial cells differentiated into AECs and SECs from 2.5 to 3.5 dpc (Fig. 6A). Therefore, we compared our dataset and the human endometrium GSE111976 dataset^[26]^ to assess their degree of correspondence. To determine the endometrial luminal epithelial cell similarity of mice and humans, we used the ssGSEA algorithm to calculate DEG set activity among the three types of epithelial cells in both mice and humans after transforming the mouse DEGs into homologous human genes. We found that human SOX9-positive epithelial cells were similar to mouse luminal “root” epithelial cells, while human ciliated epithelial cells were similar to muse AECs, and human unciliated epithelial cells were similar to mouse SECs (Fig 6B, 6C). These results suggested that conserved differentiation processes of endometrial luminal epithelial cells in humans and that they are similar to those in mice.

**Figure 6.**
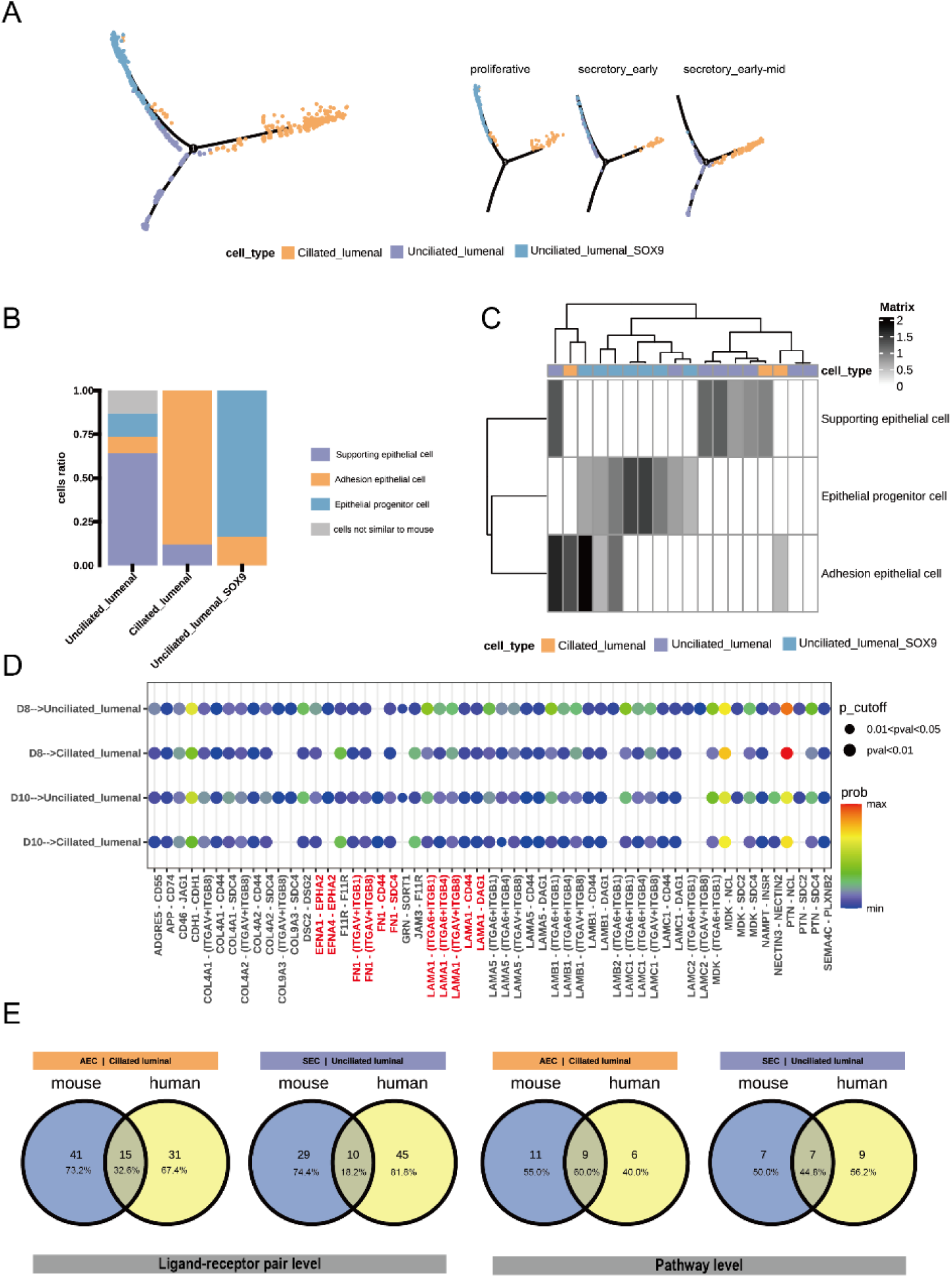
Both luminal epithelial cell differentiation and directed signaling are conserved in humans and mice. (A) Developmental pseudo-time of endometrial luminal epithelial cells in human uterus. (B) Bar plot showing predicted cell ratio of human luminal epithelial cells based on three types of mouse luminal epithelial cells. (C) Heatmap showing ssGSEA scores calculated by marker genes of three types of mouse luminal epithelial cells in human luminal epithelial cells. (D) Dot plot showing embryo –derived signals from E8.0(D8) and E10.0(D10) to ciliated luminal cells and unciliated luminal cells. Point size represented the P value and the color represented the possibility of communication of the ligand-receptor pairs between embryo and luminal epithelial cells. (E) Venn diagram showing embryo-derived signal conservation between mouse and human at ligand-receptor level (left) and pathway level(right).

As performed with mice, we also analyzed signaling crosstalk between endometrium and embryo in humans using CellChat and two human datasets, GSE109555 for the embryo and GSE111976 for the endometrium samples (Fig. 6D). To identify the signaling of embryos, we identified a total of 55 LR pairs, including previously reported FN1 and LAMA1. We found LR pairs in the EPHA pathway in humans. By analyzing the conserved communication between embryos and uteri in humans and mice, we found the signaling from embryos to ciliated luminal epithelial cells of 15 conserved LRs (32% in humans) that were involved in 9 pathways (60% in humans) and signaling from embryos to unciliated luminal epithelial cells of 10 conserved LRs (18% in humans) that were involved in 7 pathways (44.8% in humans) (Fig 6E). The LR pairs showed less species conservation than the pathways. These results indicated that in addition to the endometrial luminal epithelial cell differentiation, the signaling directing differentiation was similar between humans and mice.

### Defective proliferation and differentiation of SOX9-positive epithelial cells leads to a thin endometrium

To explore the clinical relevance of epithelial cell differentiation during peri-implantation, we used Scissor for comparing single-cell and bulk sequence datasets to predict pathology-related cell population alterations in the endometrium. First, we compared our thin endometrium bulk sequence dataset and the single-cell dataset of Hu et al^[27]^. Our bulk data indicated that terms related to the cell cycle checkpoint, negative regulation of the mitotic cell cycle and regulation of the apoptotic signaling pathway were significantly enriched (Fig. 7A, 7B), suggesting impaired cell proliferation and an excessive apoptosis rate in the thin endometrium. Based on the expression patterns in our data, the Scissor analysis predicted 1120 scissor (-) cells and 580 scissor (+) cells in Hu’s dataset (Fig. 7C). Scissor (+) cells comprised a cell population associated with the thin endometrium, and scissor (-) cells comprised a cell population associated with the normal endometrium, and the other cells were background cells and considered unrelated to this phenotype. We found that both scissor (+) and scissor (-) cells were abundantly distributed in the stromal cell population (Fig. 7D), which was consistent with the proliferation and widespread distribution of stromal cells during the proliferative phase in the endometrium. Interestingly, epithelial cells were the second most abundant cell type (Fig. 7D), with both scissor (+) and scissor (-) cells distributed in SOX9-positive epithelial cell populations (Fig. 7E, 7F). The comparison of the transcriptome between scissor (+) and scissor (-) SOX9-positive epithelial cells confirmed that processes associated with negative regulation of the cell cycle were significantly upregulated, while those associated with positive regulation of cell population proliferation were significantly downregulated. Processes associated with the negative regulation of the WNT signaling pathway were upregulated, while those associated with positive regulation of the NOTCH signaling pathway were downregulated (Fig. 7G). Notably, activation of the NOTHCH signaling pathway in the differentiation of secretary luminal and glandular epithelial cells in the endometrium had already been proven by Garcia-Alonso et al. We calculated the ssGSEA scores of the scissor (+) and scissor (-) SOX9-positive epithelial cell signatures. We found that no significant difference in the scores of the scissor (-) and control cells (P=0.39), although the scores of the control group were higher than those of the thin endometrium, and for the scissor (+) cells, the scores of thin endometrium groups were significantly higher than those of control group (P=0.026) (Fig. 7H). All these results indicated that scissor (+) and scissor (-) cells truly corresponded to thin endometrium pathogenesis and that the dysfunction of SOX9-positive epithelial cells was related to thin endometrium.

**Figure 7.**
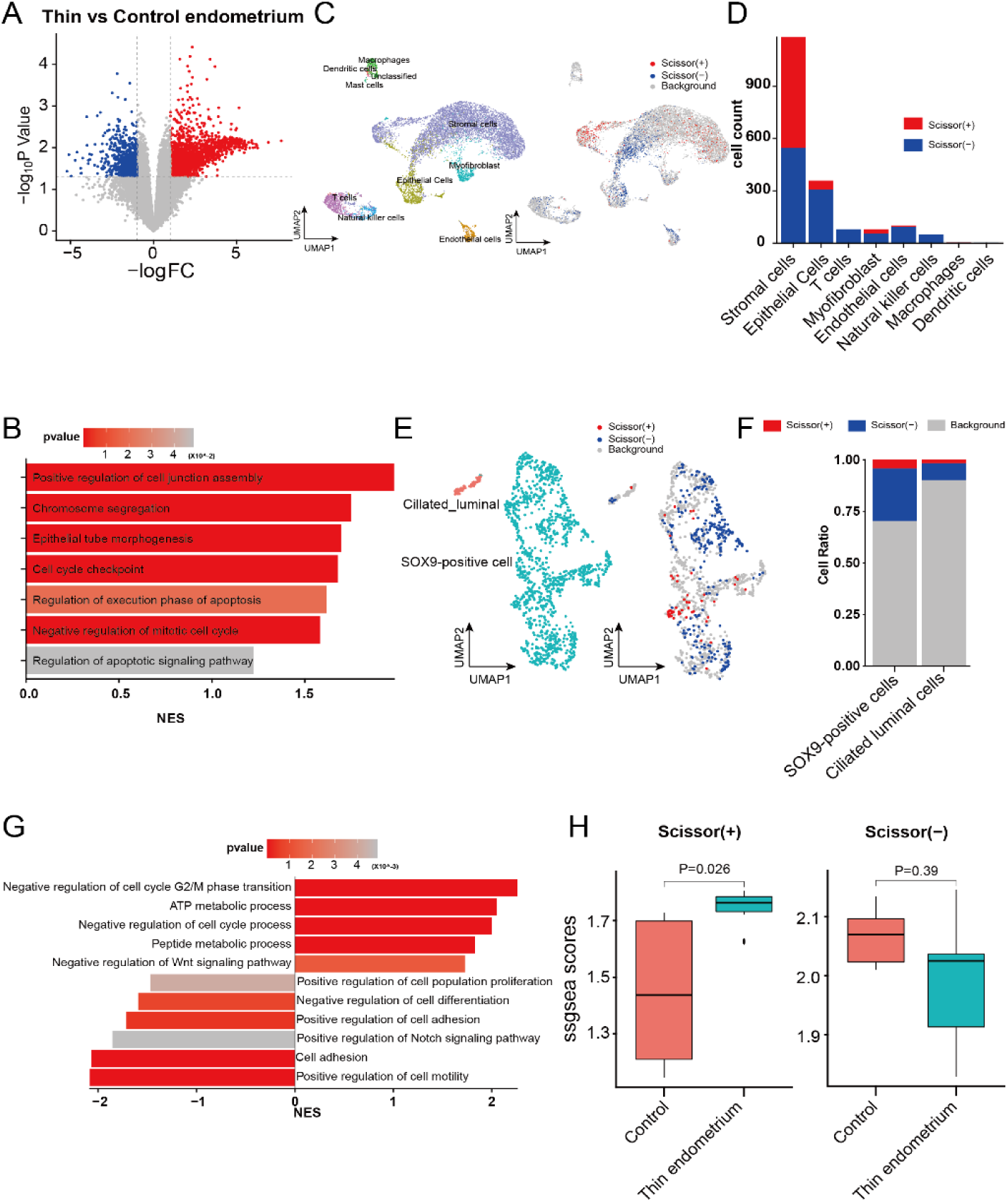
Defective proliferation and differentiation of SOX9-positive epithelial cells leads to a thin endometrium. (A) Volcano plot showing DEGs between thin and control endometrium. (B) Bar plot showing GSEA scores of enriched GO terms of DEGs found in Fig.7A. (C) UMAP showing the cell types in thin and control endometrium(left) and distribution of the cells labeled based on Scissor tool. The red and blue points at right were cells associated with thin and control endometrium phenotypes, respectively. (D) Bar plot showing the distribution of Scissor (+) and Scissor (-) cells in different cell types in human endometrium. (E) UMAP showing the cell types in luminal epithelial cells of thin and control endometrium (left) and distribution of the cells labeled based on Scissor tool. The red and blue points at right were cells associated with thin and control endometrium phenotypes, respectively. (F) Bar plot showing the distribution of Scissor (+) and Scissor (-) cells in luminal epithelial cells in human endometrium. (G) Bar plot showing ssGSEA scores of enriched GO terms of DEGs between Scissor (+) and Scissor (-) cells in SOX9-positive cells. (H) Box plot showing ssGSEA scores based on marker genes of Scissor (+) and Scissor (-) cells in control and thin endometrium.

In conclusion, our results revealed that in thin endometrium, the impaired proliferation and differentiation capacity of the luminal “root” epithelial cells affected endometrial thickness.

## Discussion

Embryo implantation requires comprehensive maternal-fetal dialog, but the temporal and cell-specific coordination of this communication is largely unknown because of the cellular complexity and dynamic developmental processes of both the embryo and endometrium during peri-implantation. By comparing the scRNA-seq data of whole uteri of pregnant and pseudo-pregnant mice when embryos were in the oviduct (2.5 dpc) and after they had entered the uterus (after 3.5 dpc) and analyzing the crosstalk indicated by bulk sequencing data of the embryos and scRNA-seq of the corresponding luminal epithelial cells in the endometrium, we revealed maternal signaling-dependent functional differentiation of luminal epithelial cells in which estrogen-responsive epithelial cells into two new types of epithelial cells, adhesion epithelial cells (AECs) and supporting epithelial cells (SECs) during 2.5 to 3.5 dpc, which initiated receptivity in conjunction with other endometrial cell types, such as stromal cells. In addition to maternal estrogen and progesterone signaling, embryonic signaling further activated AEC and SEC activity, enhancing the attachment of embryos to the endometrium. Furthermore, embryonic signaling induced additional transformation of the AECs by preventing adjacent AECs, but not AECs away from the embryo in the implantation chamber, from undergoing apoptosis. Our work provides comprehensive and systematic information on endometrial luminal epithelial cell development directed by maternal and embryonic signaling, which is important for receptivity establishment and embryo implantation.

During receptivity establishment, the hormone-induced switch in endometrial cell activity is an indicator that the uterus is ready for implantation of an embryo entering the uterus^[28-30]^. Garcia-Alonso et al^[26]^ reported that the human epithelium in the secretory phase could be divided into secretory epithelium (glandular) and ciliated epithelium (luminal), but the functional differentiation of the luminal epithelial cells caused by hormones had not been reported. In this study, we found that on 2.5 dpc, a group of estrogen-responsive luminal epithelial cells were generated by estrogen stimulation in the mouse endometrium, and then, these epithelial cells differentiated into two groups of functional cells, AECs and SECs, in response to progesterone but not embryo-derived signaling. It has been reported that mice with Pgr specifically knocked out in uterine epithelial cells showed defective epithelial cell proliferation and embryo adhesion failure. We have shown that progesterone regulated the differentiation of the uterine epithelium by activating large amounts of TFs in uterine luminal epithelium cells mediated through Pgr; these TFs included Twist1 in AECs and Fosl2 in SECs, which control switching between uterine epithelial cell proliferation and differentiation of and affects uterine receptivity^[31][32-34]^. Moreover, our work is corroborated by a recent study of human endometrial epithelial cell differentiation. The SOX9^+^/LGR5^+^ epithelial cells in the recent study constituted a population that maintained a highly similar gene characteristic that was similar to that in the luminal epithelial cells in our study, and these cells showed the potential to differentiate into unciliated and ciliated epithelial cells. Thus, the luminal epithelial cell differentiation we described during receptivity establishment in the mouse uterus was highly conserved among species. More importantly, we found that in thin endometrium, in addition to that of stromal cells, the function of luminal epithelial cells was impaired. epithelial cells differentiation-related signaling^[26]^, such as WNT and NOTCH signaling, was downregulated in thin endometrium-related scissor (+) luminal epithelial cells. These results indicated that the impaired proliferation and differentiation in the luminal epithelium led to thin endometrium.

Embryo implantation is a critical event after receptivity is established and relies on proper maternal-embryonic crosstalk. Considering on the aforementioned results, we found that changes in the transcriptome in the mouse AECs on 4.0 dpc were more dramatic than those in mice on day 4.0 of pseudo-pregnancy, which indicated that AEC activity was specifically regulated by embryo-derived signaling but not maternal signaling. Because we were limited to bulk RNA-seq data of the endometrium obtained in previous studies, it was difficult to precisely determine the crosstalk of the embryo and uterus. In our study, we resolved many signals transmitted from the embryo to AECs and SECs, such as WNT, SPP1, PDGF and EPHA signals at single-cell resolution. It has been previously shown that Ddk1-mediated the inhibition of WNT/β-catenin signaling, which had no negative effect on mouse uterine receptivity but significantly inhibited embryo adhesion^[35]^. Moreover, SPP1 signaling promotes embryo adhesion by binding ITGAV^[36]^. In the present study, we established a more systematic and comprehensive regulatory network of embryo-derived signaling, especially signaling in two luminal epithelial cell subpopulations: AECs and SECs. Furthermore, we found that the functional effects of these signaling pathways on AECs and SECs were related to epithelial cell remodeling processes and protein phosphorylation, respectively, which may be consistent with the adhesion function of AECs and the nutritional and supportive function of SECs. Interestingly, we found that on 4.5 dpc, AECs were mainly distributed near the embryo implantation site, although they were also distributed in the full lumen on 4.0 dpc, which may have been because embryo-derived signaling prevented these cells from undergoing apoptosis. This finding further illustrated the effect of embryo-derived signaling on the spatial distribution of the endometrial epithelial cells. Furthermore, we demonstrate the promoting function of the key embryo-derived signaling of Pdgfa and Efna3/4 in embryo implantation. According to previously obtained transcriptome data from trophoblast cells and endometrial cells during embryo implantation, Pdgfa expression was upregulated in human trophoblast cells, and its corresponding receptor, Pdgfra, was overexpressed in the endometrium of pregnant women^[9]^, suggesting that PDGF signaling is involved in the early dialog between blastocysts and maternal endometrial cells. On the other hand, Epha1 was mainly expressed in the mouse endometrial epithelial cells, while its ligand ephrin A1– 4 was expressed in blastocysts^[37]^. In our results, the inhibition of PDGF signaling significantly reduced embryo implantation efficiency. Similarly, after we targeted siRNAs to the expression of Epha1 in the uterus, the embryo implantation efficiency also decreased significantly. These results indicated that PDGF and EPHA signaling of the embryo acted on the luminal epithelium, playing an important role in embryo implantation. Furthermore, comparing our data with single-cell data obtained from previously obtained human receptive endometrium samples^[25]^, we found that maternal-embryo crosstalk involving CDH, COL1A2, FGF, SPP1, and WNT was highly conserved among species.

In conclusion, our results revealed the dynamics and molecular mechanisms of the epithelium during the establishment of receptivity and embryo implantation under the regulation of maternal and embryonic signaling and provide a reference for understanding the cytological behavior during the establishment of endometrial receptivity. Specifically, by determining the degree of conservation of epithelial cell differentiation and maternal-embryo communication among humans and mice, we demonstrated the association of impaired epithelial cell differentiation with endometrial function in thin endometrium.

## Acknowledgements

This work was supported by National Natural Science Foundation of China (82030040, 81871165, 81871165, 31872846, 8197138, 81901444, 82001459). We thank OE Biotech Co., Ltd (Shanghai, China) and LC-BIO technologies CO., Ltd ((Hangzhou)) for providing single-cell RNA-seq and bulk RNA-seq.

## Supplemental Figures

**Supplemental figure 1.**
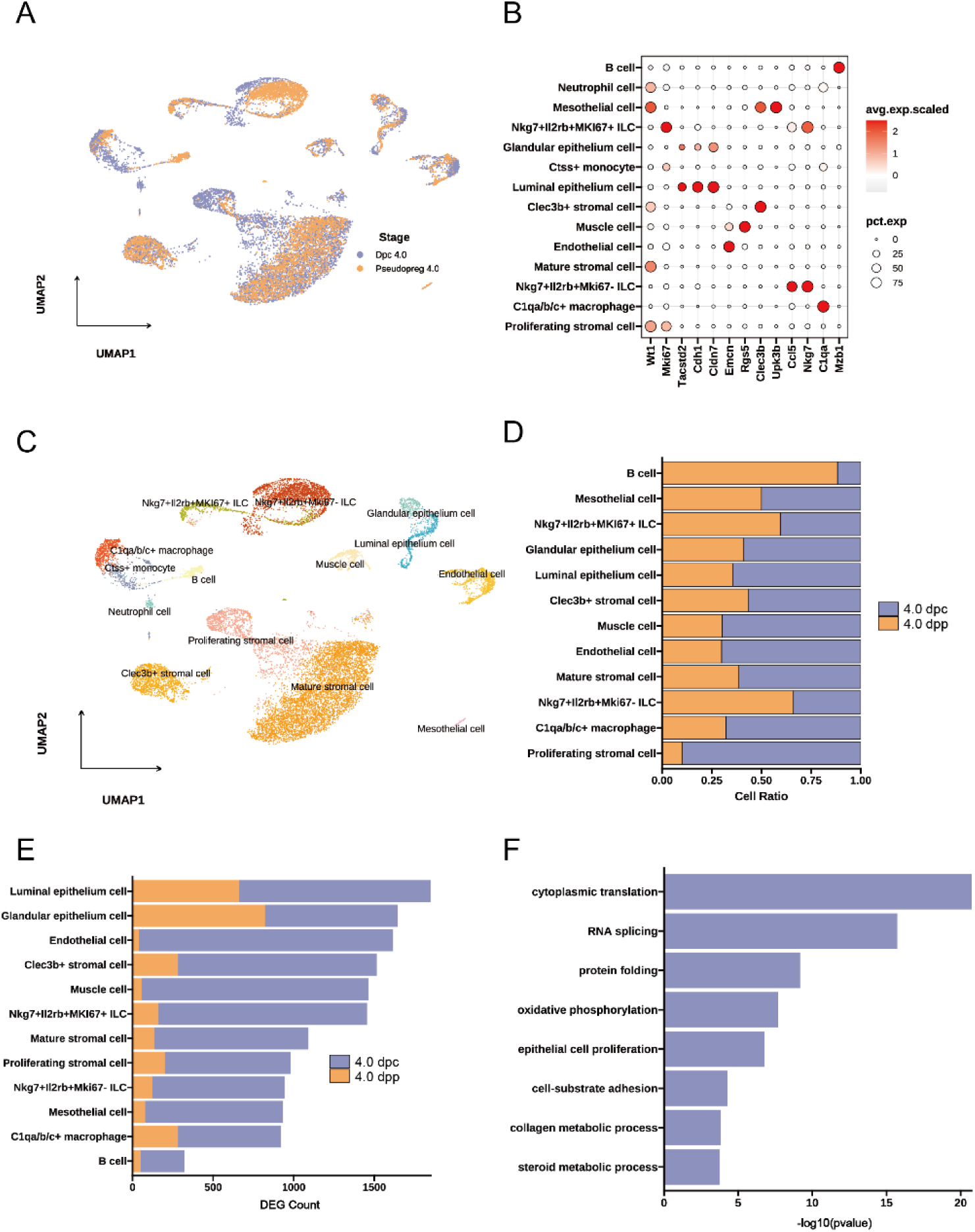
Endometrial luminal epithelial cells show dramatic transcriptomic changes in the uterine receptivity state. (A) UMAP projections of scRNA-seq data from uteri of mice on 4.0 dpc and 4.0 dpp. (B) Dot plot showing log2-transformed expression of marker genes to identity cell types in uteri of mice on 4.0 dpc and 4.0 dpp. (C) UMAP showing cell types from uteri of mice on 4.0 dpc and 4.0 dpp. (D) Bar plot showing relative proportion of cells in uteri of mice on 4.0 dpc and 4.0 dpp. (E) Bar plot showing the count of differential expressed genes in endometrial cell population between uteri of mice on 4.0 dpc and 4.0 dpp. (F) Bar plot showing enriched GO terms of up-regulated in luminal epithelial cells of mice at 4.0 dpc.

**Supplemental figure 2.**
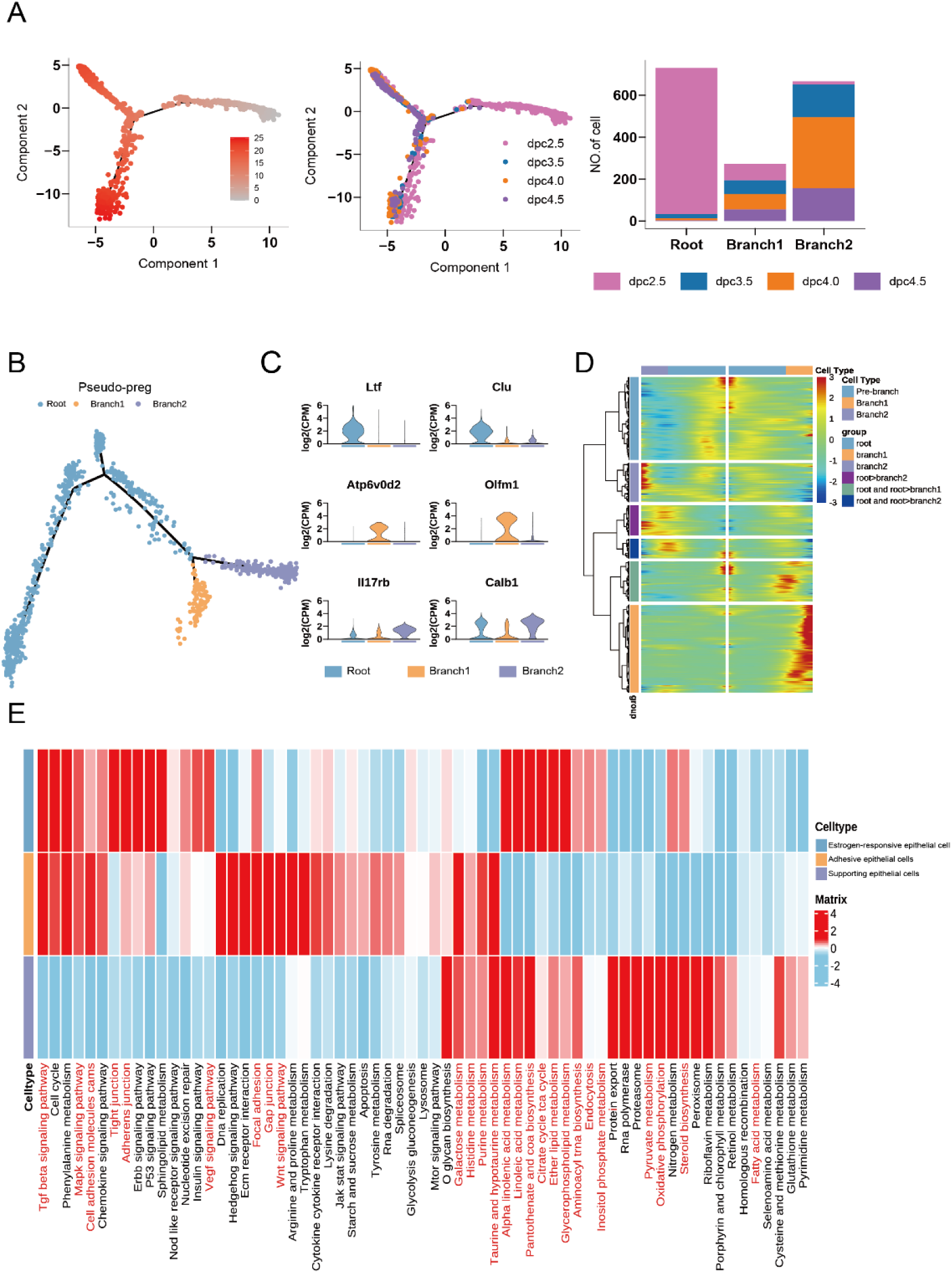
Pseudo-time analysis of endometrial luminal epithelial cells in pregnant and pseudo0pregnant mice. (A) Dot plot showing cell trajectories ordered in pseudo-time(left) and time points(middle) and bar plot showing the distribution of the three cell types in each time point. (B) Developmental pseudo-time of endometrial luminal epithelial cells in pseudo-pregnant mice. (C) Violin plot showing log2-transformed expression of marker genes in three types of luminal epithelial cells in mice. (D) Heatmap showing hierarchical relationship between clusters of genes that were differentially expressed across pseudo-time from endometrium in pregnant mice. (E) Heatmap showing GSVA scores of KEGG pathways in all three types of luminal epithelial cells in mice.

**Supplemental figure 3.**
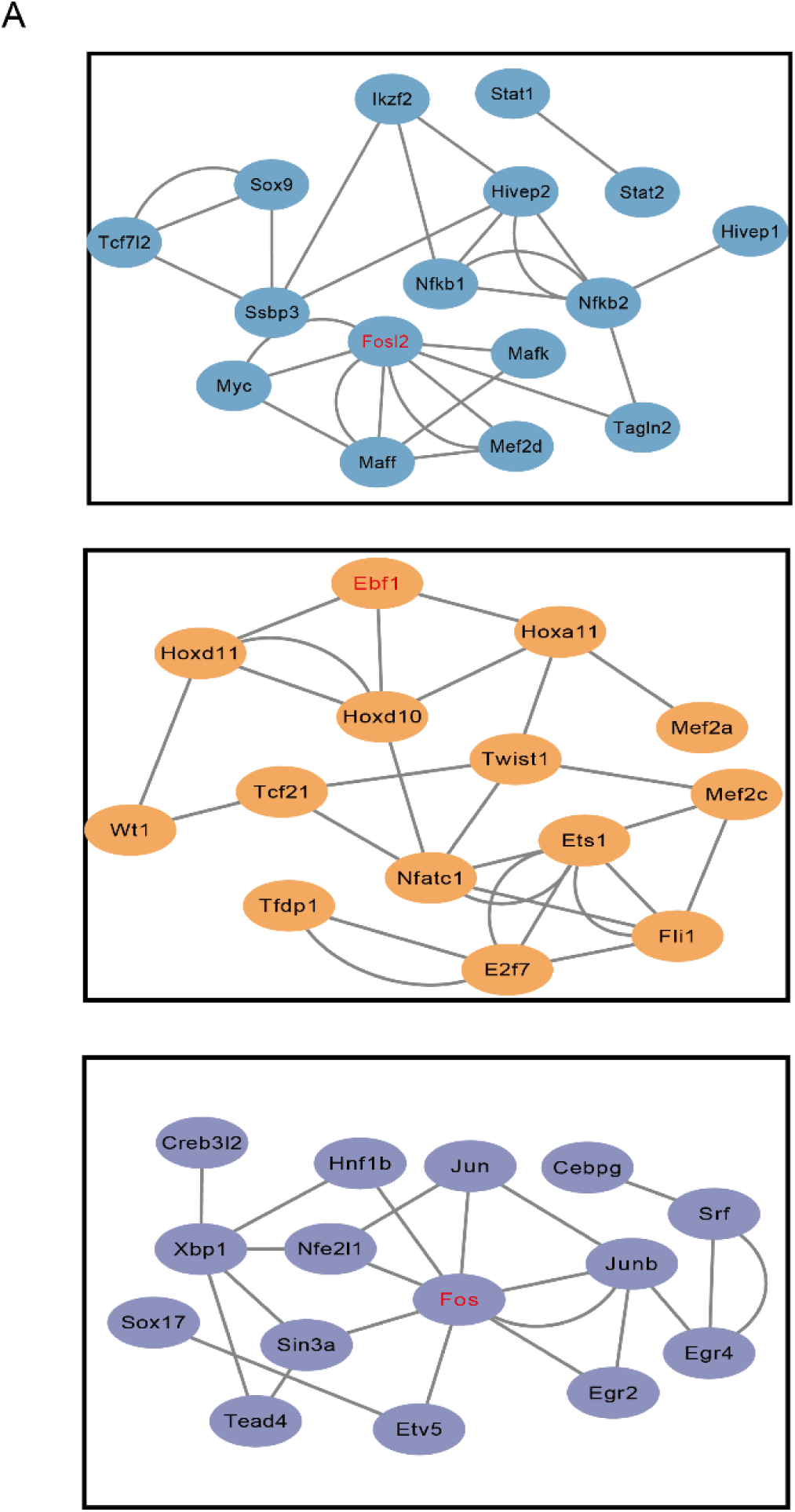
Regulatory networks of transcription factors during luminal epithelial cell differentiation. (A) Regulatory networks of transcription factors during luminal epithelial cell differentiation in Root (top), AECs (middle) and SECs (bottom).

## Reference

[1] Gellersen B, Brosens JJ. Cyclic decidualization of the human endometrium in reproductive health and failure. Endocr Rev, 2014, 35: 851–905

[2] Jabbour HN, Kelly RW, Fraser HM, et al. Endocrine regulation of menstruation. Endocr Rev, 2006, 27: 17–46

[3] Hertz-Picciotto I, Samuels SJ. Incidence of early loss of pregnancy. N Engl J Med, 1988, 319: 1483–1484

[4] Kim SM, Kim JS. A review of mechanisms of implantation. Dev Reprod, 2017, 21: 351–359

[5] Lessey BA. Assessment of endometrial receptivity. Fertil Steril, 2011, 96: 522–529

[6] Cha J, Sun X, Dey SK. Mechanisms of implantation: Strategies for successful pregnancy. Nat Med, 2012, 18: 1754–1767

[7] Paulson RJ. Hormonal induction of endometrial receptivity. Fertil Steril, 2011, 96: 530–535

[8] Peter Durairaj RR, Aberkane A, Polanski L, et al. Deregulation of the endometrial stromal cell secretome precedes embryo implantation failure. Mol Hum Reprod, 2017, 23: 582

[9] Haouzi D, Dechaud H, Assou S, et al. Transcriptome analysis reveals dialogues between human trophectoderm and endometrial cells during the implantation period. Hum Reprod, 2011, 26: 1440–1449

[10] Tapia-Pizarro A, Archiles S, Argandona F, et al. Hcg activates epac-erk1/2 signaling regulating progesterone receptor expression and function in human endometrial stromal cells. Mol Hum Reprod, 2017, 23: 393–405

[11] Paria BC, Ma W, Tan J, et al. Cellular and molecular responses of the uterus to embryo implantation can be elicited by locally applied growth factors. Proc Natl Acad Sci U S A, 2001, 98: 1047–1052

[12] Hamatani T, Daikoku T, Wang H, et al. Global gene expression analysis identifies molecular pathways distinguishing blastocyst dormancy and activation. Proc Natl Acad Sci U S A, 2004, 101: 10326–10331

[13] Maccarrone M, DeFelici M, Klinger FG, et al. Mouse blastocysts release a lipid which activates anandamide hydrolase in intact uterus. Mol Hum Reprod, 2004, 10: 215–221

[14] Piao HL, Tao Y, Zhu R, et al. The cxcl12/cxcr4 axis is involved in the maintenance of th2 bias at the maternal/fetal interface in early human pregnancy. Cell Mol Immunol, 2012, 9: 423–430

[15] Svensson V, Vento-Tormo R, Teichmann SA. Exponential scaling of single-cell rna-seq in the past decade. Nat Protoc, 2018, 13: 599–604

[16] Hao Y, Hao S, Andersen-Nissen E, et al. Integrated analysis of multimodal single-cell data. Cell, 2021, 184: 3573–3587 e3529

[17] Korsunsky I, Millard N, Fan J, et al. Fast, sensitive and accurate integration of single-cell data with harmony. Nat Methods, 2019, 16: 1289–1296

[18] Trapnell C, Cacchiarelli D, Grimsby J, et al. The dynamics and regulators of cell fate decisions are revealed by pseudotemporal ordering of single cells. Nat Biotechnol, 2014, 32: 381–386

[19] Aibar S, Gonzalez-Blas CB, Moerman T, et al. Scenic: Single-cell regulatory network inference and clustering. Nat Methods, 2017, 14: 1083–1086

[20] Jin S, Guerrero-Juarez CF, Zhang L, et al. Inference and analysis of cell-cell communication using cellchat. Nat Commun, 2021, 12: 1088

[21] Wu T, Hu E, Xu S, et al. Clusterprofiler 4.0: A universal enrichment tool for interpreting omics data. Innovation (N Y), 2021, 2: 100141

[22] Hanzelmann S, Castelo R, Guinney J. Gsva: Gene set variation analysis for microarray and rna-seq data. BMC Bioinformatics, 2013, 14: 7

[23] Blum A, Mostow K, Jackett K, et al. Klf4 regulates metabolic homeostasis in response to stress. Cells, 2021, 10:

[24] Rubel CA, Lanz RB, Kommagani R, et al. Research resource: Genome-wide profiling of progesterone receptor binding in the mouse uterus. Mol Endocrinol, 2012, 26: 1428–1442

[25] Wang W, Vilella F, Alama P, et al. Single-cell transcriptomic atlas of the human endometrium during the menstrual cycle. Nat Med, 2020,

[26] Garcia-Alonso L, Handfield LF, Roberts K, et al. Mapping the temporal and spatial dynamics of the human endometrium in vivo and in vitro. Nat Genet, 2021, 53: 1698–1711

[27] Lv H, Zhao G, Jiang P, et al. Deciphering the endometrial niche of human thin endometrium at single-cell resolution. Proc Natl Acad Sci U S A, 2022, 119:

[28] Hirota Y. Progesterone governs endometrial proliferation-differentiation switching and blastocyst implantation. Endocr J, 2019, 66: 199–206

[29] Dey SK, Lim H, Das SK, et al. Molecular cues to implantation. Endocr Rev, 2004, 25: 341–373

[30] Haraguchi H, Saito-Fujita T, Hirota Y, et al. Microrna-200a locally attenuates progesterone signaling in the cervix, preventing embryo implantation. Mol Endocrinol, 2014, 28: 1108–1117

[31] Fukui Y, Hirota Y, Matsuo M, et al. Uterine receptivity, embryo attachment, and embryo invasion: Multistep processes in embryo implantation. Reproductive medicine and biology, 2019, 18: 234–240

[32] Lee K, Jeong J, Kwak I, et al. Indian hedgehog is a major mediator of progesterone signaling in the mouse uterus. Nat Genet, 2006, 38: 1204–1209

[33] Li Q, Kannan A, DeMayo FJ, et al. The antiproliferative action of progesterone in uterine epithelium is mediated by hand2. Science, 2011, 331: 912–916

[34] Xin Q, Kong S, Yan J, et al. Polycomb subunit bmi1 determines uterine progesterone responsiveness essential for normal embryo implantation. J Clin Invest, 2018, 128: 175–189

[35] Xie H, Tranguch S, Jia X, et al. Inactivation of nuclear wnt-beta-catenin signaling limits blastocyst competency for implantation. Development, 2008, 135: 717–727

[36] Frank JW, Seo H, Burghardt RC, et al. Itgav (alpha v integrins) bind spp1 (osteopontin) to support trophoblast cell adhesion. Reproduction (Cambridge, England), 2017, 153: 695–706

[37] Fujii H, Tatsumi K, Kosaka K, et al. Eph-ephrin a system regulates murine blastocyst attachment and spreading. Dev Dyn, 2006, 235: 3250–3258

